# Establishment of a colony of *Anopheles darlingi* from French Guiana for vector competence studies on malaria transmission

**DOI:** 10.1101/2022.05.24.493327

**Authors:** Nicolas Puchot, Marie-Thérèse Lecoq, Romuald Carinci, Jean Bernard Duchemin, Mathilde Gendrin, Catherine Bourgouin

**Author notes:** **Correspondence:** Catherine Bourgouin.

## Abstract

*Anopheles darlingi* is a major vector of both *Plasmodium falciparum* and *Plasmodium vivax* in South and Central America. However, vector competence and physiology of this mosquito species have been scarcely studied due to difficulties in rearing it in the laboratory. Here, we report the successful establishment of a robust colony, from mosquito collection in French Guiana. We describe our mosquito colonization procedure with relevant information on environmental conditions, mating ability, larval development and survival, recorded over the first six critical generations. Experimental infection showed that our *An. darlingi* colony has a moderate permissiveness to *in vitro* produced gametocytes of the *P. falciparum* NF54 strain originating from Africa. This colony, that has reached its 20^th^ generation, will allow further characterization of *An. darlingi* life-history traits and of *Plasmodium*-mosquito interactions with South American malaria parasites.

## Introduction

The establishment of mosquito colonies has been instrumental for a better understanding of many of their biological features including development, insecticide resistance and pathogen transmission. It is also important to test novel insecticides and novel technologies for population suppression or replacement such as the sterile insect technique, the RIDL approach (Release of Insects carrying Dominant Lethals), the use of symbiont bacteria such as *Asaia* or the production of transgenic mosquitoes.

Among mosquito species known as vectors of human pathogens, species belonging to the *Anopheles* genus and vector of malaria parasites are the most numerous and have the most widespread distribution [1]. In addition, *Anopheles* species have specific reproductive characteristics notably their eurygamous behavior i.e. their propensity to mate in swarms and the female refractoriness to multiple mating. Those characteristics have been challenging for successful establishment of colonies of many *Anopheles* species. To overcome these limitations, one strategy is the forced mating technique, which relies on the manual adjustment of male and female copulatory structures [2–4]. Although tricky and highly demanding in human resources, this technique was successful for maintaining some species as laboratory colony for many years. The methodology has notably been used to colonize and maintain *Anopheles dirus* and is still used to create and maintain novel *Anopheles* colonies [5]. Interestingly, some species such as *Anopheles albitarsis* were reported to change to stenogamous free-mating behaviour after several generations of colony maintenance by forced copulation [6].

*Anopheles darlingi* (also called *Nyssorhynchus darlingi* [7]) is a major vector of both *Plasmodium falciparum* and *Plasmodium vivax* in South and Central America [8]. As most anophelines, *An. darlingi* has an eurygamous behaviour that precludes its readiness to mate in captivity. In contrast to many *Anopheles* species belonging to the *Cellia* and *Anopheles* subgenera, it is not clear whether *Anopheles* belonging to the *Nyssorhynchus* subgenus exhibit a swarming behavior for mating [9]. Indeed, Lounibos *et al*. reported that young F1 virgin *An. darlingi* females would mate within 2 to 36 h after being released in a village where no mosquito swarm was evidenced [10]. In 61 cm-high cages, Villarreal-Treviño *et al*. observed pseudo-swarms of *An. darlingi* occurring 5-40cm above the cage floor [11]. This may be due to space constraints yet corroborates field observations that males fly at low altitude. As *An. darlingi* is a major malaria vector in South America and particularly in the Amazon basin, several teams tried to colonize this mosquito species for vector competence studies and possibly human *Plasmodium* sporozoite production. The oldest published records of the establishment of a free-mating *An. darlingi* colony go back to 1947 with *An. darlingi* from the Co-operative Republic of Guyana (at that time British Guiana), Colombia and Brazil [12–14]. Since then, two reports of Brazilian (Araraquara, Sao Paulo State) *An. darlingi* colonies were published in 1970 and 1988 [15, 16] and followed almost 40 years later with the description of three novel free-mating *An. darlingi* colonies, two from Peru and one from the Rondonia State (Amazon basin – Brazil) [11, 17, 18].

Here, we described the establishment of a free-mating colony of *An. darlingi* from a different South America region, French Guiana. By mixing our expertise in *Anopheles* mosquitoes to published results and improvements along the most recent established *Anopheles pseudopunctipennis* and *An. darlingi* free-mating colonies [11, 17, 19, 20], this novel *An. darlingi* colony was produced in less than 8 months and has currently reached its 20^th^ generation.

## Material and Methods

### Producing the F1 generation from wild Anopheles darlingi

Wild *An. darlingi* were captured in Val’ranch (Tonate Macouria village; 05.01616°N 052.52845W), a location known for its abundance in this species [21], using two Mosquito Magnet® MM300 Executive traps arranged 50 m apart. The capture was performed over a single night with two catching sessions from 6 pm to 8 pm, and from 8 pm to 7 am in September 2020. Mosquito females were transferred from the traps into 30 cm cubic plastic cages (Bugdorm®) covered with damp towels to maintain high humidity. They were transported to the laboratory (Institut Pasteur de la Guyane, Cayenne, French Guiana) in a foam box to buffer ambient temperature. There, they were fed on anesthetized mice. On the fourth day after feeding, gravid females to which one wing was removed were transferred to individual Eppendorf tubes containing wet filter paper as a forced-egg laying method [22]. Once the eggs were laid, the females were removed before air freight transfer of tubes with egg clutches to Institut Pasteur (Paris, France).

### Insectary parameters

Following the publications on *An. pseudopunctipennis* [19] and *An. darlingi* [11, 17], the light periodicity was set up at 12:12 with dawn and dusk spanning each 1h. To stimulate copulation, blue light flashes (Led stroboscopic light) were provided for 45 min just after full darkness at a rate of 140 flashes per minute, each flash lasting ~0.2 s. Temperature in French Guiana cycles daily between 22 and 32°C when *An. darlingi* can be found (Suppl Fig. 1 and Suppl file; https://donneespubliques.meteofrance.fr/?fond=produit&id_produit=90&id_rubrique=32-acces date 2019/08/23). Considering these natural temperature conditions and expecting an adapted behavior keeping mosquitoes away from high temperatures in the field, the insectary temperature was set at 26°C during the daytime and down to 23°C during the night time. A constant relative humidity was kept around 75%, which is close to the average humidity observed in the field during *An. darlingi* peaks (81%, Suppl Fig. 1). A summary of those experimental conditions is presented in Fig. 1.

**Figure 1:**
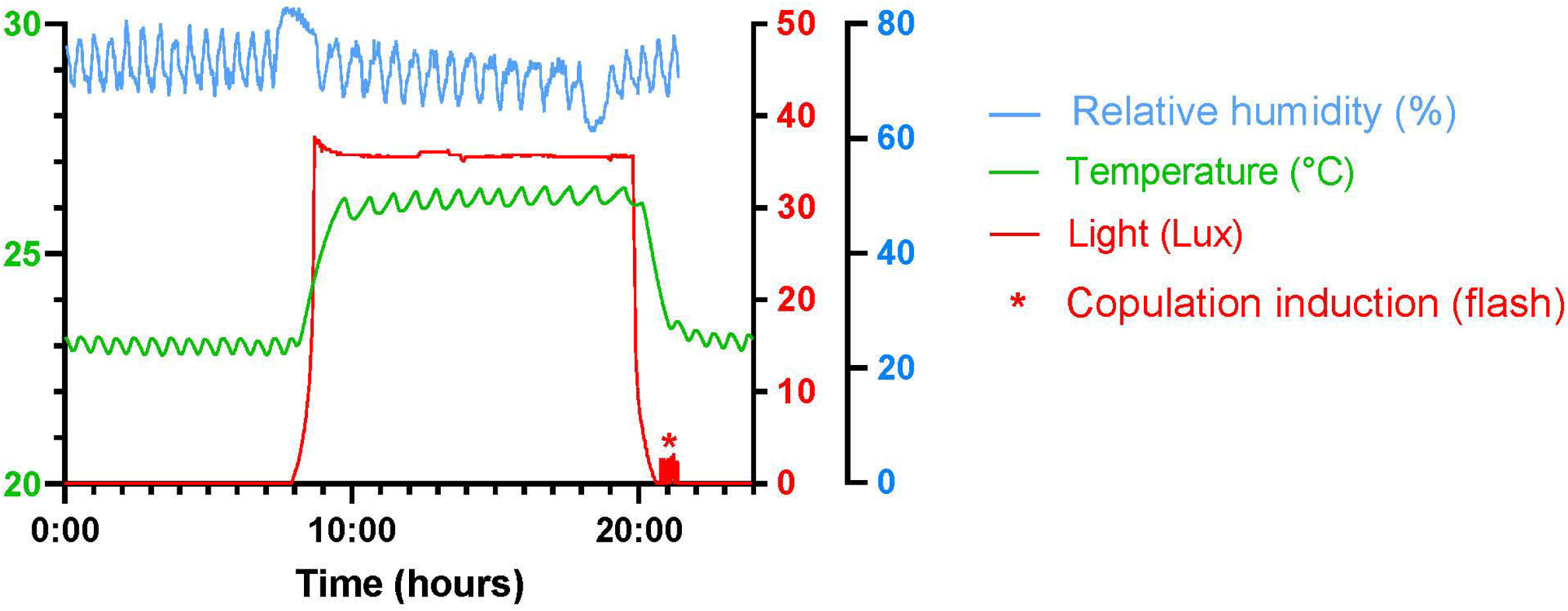
Insectary parameters. The graph represents temperature (green), relative humidity (blue) and light intensity (red) recorded via a Hobo 1019 recorder (MX1104). Temperature and humidity parameters are provided by a dedicated automated system. Light (led) is controlled by an electronic device. Blue light flashes are provided by a led stroboscopic light, just after dusk (red asterisk).

### Mosquito rearing

Larvae were reared in white square plastic pans filled to 2 cm height with demineralized water supplemented with a mixture of salts (200X solution: NaCl 0.2%; CaSO_4-_2H_2_O 0.3%; NaHCO_3_ 1%) and covered with a mosquito net. L1 to L2 larvae were fed with Tetra®TetraMin Baby and later stages with cat food. When the first pupae appeared, a small container filled with a sterile 10% sucrose solution in mineral water (Volvic®) and holding a dental cotton roll (ROEKO Parotisroll Coltene-10 cm height) was placed in the center of the rearing tray to feed emerging adults. Adults were collected with an in-house made electric aspirator and transferred in mating cages (47.5×47.5×47.5 cm; BugDorm-4F4545 - https://shop.bugdorm.com). Because of the size of the cage, two sucrose solution containers were placed at different heights in the cage. They were replaced every other day.

At each generation, mosquitoes were given 5 to 7 days for free mating (with light-flash induction, as described above) before being fed on anesthetized animals, either rabbit or mice (See details in the Results).

Females were allowed to lay eggs in a black plastic tray filled to 1.5 cm height with a 2:1 mix of demineralized water and filter-sterilized water (22μm filters) from the larval containers as a possible attractive cue to stimulate females to deposit their eggs. Eggs were collected by filtration through an in-house made strainer equipped with Nitex (125 μm fine-mesh - Dutscher #987452) and then transferred for hatching in a rearing pan.

### Experimental infections with Plasmodium berghei and Plasmodium falciparum

Five-day-old *An. darlingi* females were infected with *P. berghei* (GFP ANKA@HSP70 clone) or *P. falciparum* NF54, concurrently with same age *An. stephensi* females (Sda500 strain). The mosquito infections were performed by the local *Anopheles* infection platform (CEPIA, Institut Pasteur). Before being exposed to an infected blood meal, females were starved from sugar overnight with access to water only in the form of a damp cotton ball, which was removed 4 h before blood feeding. Mosquito infection with *P. berghei* was performed by placing 2 anesthetized infected mice on the cage for 15 min. Only fully fed females were maintained for 10 days at 21 °C for oocyst development. Mosquito infection with *P. falciparum* was performed by standard membrane feeding assay (SMFA) using glass feeders containing *in vitro* produced gametocytes as described previously [23]. Fully fed females were transferred to small cages and maintained for 7 days (oocyst detection) at 26°C.

Oocysts were detected on dissected midguts stained with 0.1% mercuro-bromo fluorescein (Fluka) in 1X phosphate buffered saline.

### Animals- Ethic Statement

This study was conducted in strict accordance with the recommendations from the Guide for the Care and Use of Laboratory Animals of the European Union (European Directive 2010/63/UE) and the French Government. The protocol for Institut Pasteur-Paris was approved by the “Comité d’éthique en expérimentation animale de l’Institut Pasteur” CETEA 89 (Permit number: 2013–0132), by the French Ministry of Scientific Research (Permit number: 02174.02). For Institut Pasteur de La Guyane the protocol was approved by the French Ministry of Scientific Research (Permit number APAFIS 2016081011367627_v5). Mosquito field collection in French Guiana were performed according to the Nagoya Protocol (Number 388019 - 19 March 2019).

## Results

### Establishing the first 6 generations of free-mating An. darlingi

From 34 wild *An. darlingi* females, which had successfully blood fed and laid eggs in tubes, we obtained ~2,000 F1 eggs that led to 705 adults with a 1:1 sex ratio. Seven days after the first F1 adults emerged, mosquitoes were fed on anesthetized animals on 9 consecutive days followed by 4 additional feedings spaced roughly every three days (Fig. 2). To avoid mosquito drowning in the water of the egg laying container, this was provided to females after the fifth blood meal only. The container was inspected every day for egg laying and the first eggs were laid after the 6^th^ blood meal. Only 20 eggs were collected on that day. Although eggs were collected almost every day since the 6^th^ blood meal, the size of each egg clutch rarely exceeded 50 eggs. Nevertheless, combining the frequency of blood feedings and egg collection over nearly 3 weeks allowed us to collect ~1,050 F2 eggs that produced 505 adults.

**Figure 2:**
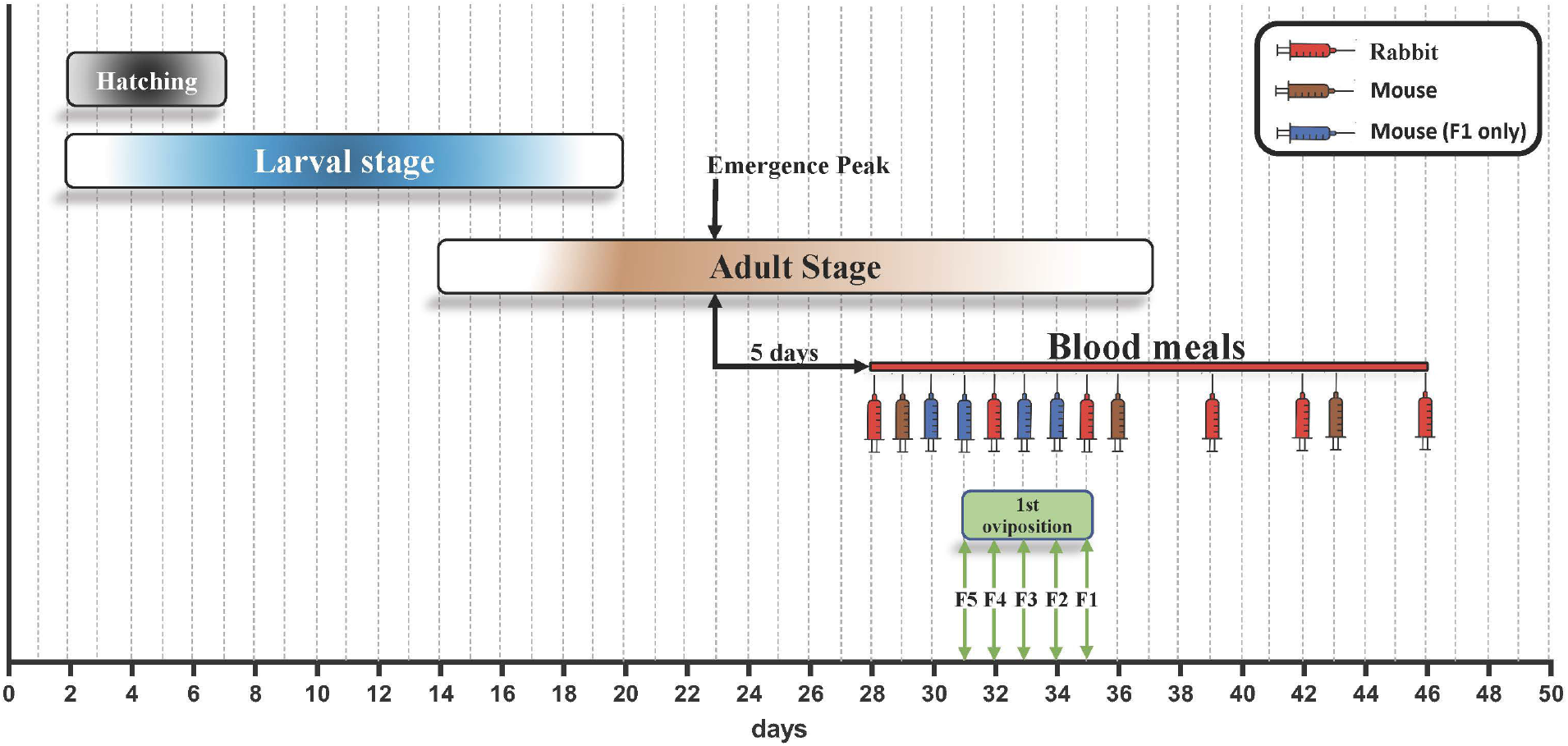
Schematic representation of the production of the first 6 generations of *An. darlingi*. The duration of egg hatching, larval stage and adult stage are colour coded. The red line is a schematic representation of the blood feeding strategy that was used over the first 6 generations by alternate feeding on rabbit (red) and mice (brown). Some additional blood feedings on mice were only provided to the first generation (blue). Between F2 and F6, the number of feedings varied based on the availability of the animals and the egg production and are detailed in Figure 3. At each generation, the time required for the females to lay the first egg batch (green) diminished across generations.

The same procedure and blood feeding scheme (Fig. 2) allowed to reach a F6 population of nearly 3,300 adults without any trouble (Fig. 3).

**Figure 3:**
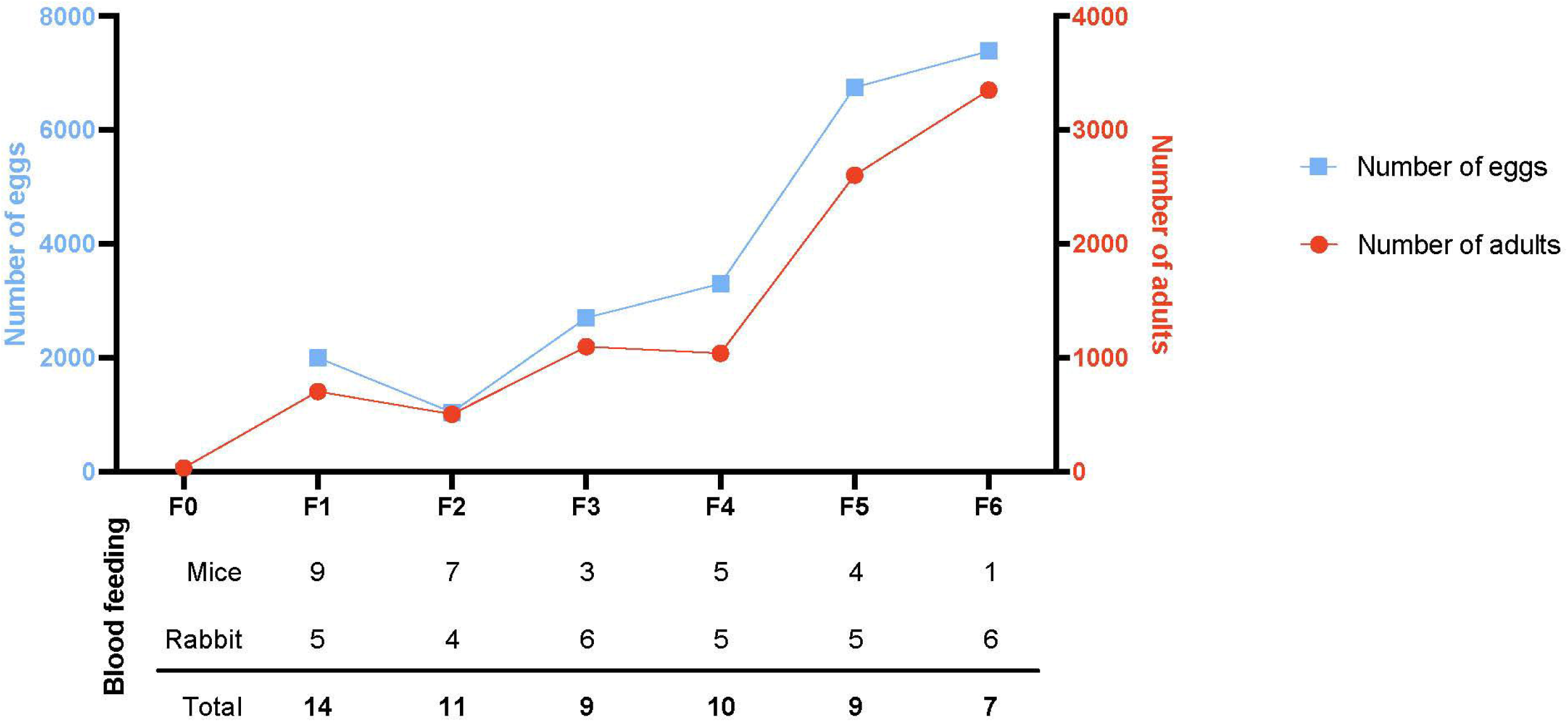
Yield in eggs and adults from F1 to F6. The graph represents the number of eggs obtained at each generation (blue line with squares and left scale) and the subsequent number of adults (red curve with circles and right scale). The number of blood feedings provided at each generation is indicated below.

### Bionomic features

#### Egg production and development

The wild caught females produced a mean of 59 eggs per blood fed female (Table 1). This number dropped between an estimated 3 to 13 eggs per female during the establishment of the colony despite providing over 9 blood meals to the females (Table 1 and Sup data S1). This low number likely represents the low number of fertilized females and not the intrinsic fecundity of fertilized females. This could be confirmed by additional investigation as spermatheca dissection and single female egg collection.

**Table 1:** This table summarizes the raw data obtained across 7 free mating generations of *An. darlingi*, starting with 34 wild caught gravid females producing 2000 F1eggs that led to the production of 705 F1 adults.

| Parental Generation | Number of eggs | Number of larvae | Numbers of females | Numbers of adults | Number of blood meals | egg eclosion rate (%) | Larvae to adults mortality (%) | Eggs /female | Eggs/female/ blood meal |
| --- | --- | --- | --- | --- | --- | --- | --- | --- | --- |
| F0 |  |  | 34 | 34 | 1 |  |  | 58.8 | 58.8 |
| F1 | 2000 | 800 | 344 | 705 | 14 | 40 | 11.88 | 3.0 | 0.2 |
| F2 | 1045 | 530 | 247 | 505 | 11 | 51 | 4.72 | 11.0 | 1.0 |
| F3 | 2705 | 1425 | 590 | 1099 | 9 | 53 | 22.88 | 5.6 | 0.6 |
| F4 | 3300 | 1780 | 578 | 1040 | 10 | 54 | 41.57 | 11.7 | 1.2 |
| F5 | 6750 | 3029 | ND | 2605 | 9 | 45 | 14.00 | 5.4 | 0.6 |
| F6 | 7390 | 3862 | ND | 3351 | 7 | 52 | 13.23 | 5.9 | 0.8 |
| F7 | 9650 | ND | ND | 4097 | 7 | ND | ND | 8.7 | 1.2 |

Eggs laid on the same day after a blood meal hatched over 2 to 7 days leading to larval stages overlapping from L1 to L3. The hatching rate was relatively stable and rarely exceeded 50% (Table 1 and Fig. 4), suggesting that unfertilized blood-fed females can lay unfertilized eggs or that the number of spermatozoids transferred to female during free-mating is not sufficient to fertilize all the oocytes developing in a single blood-fed female.

**Figure 4:**
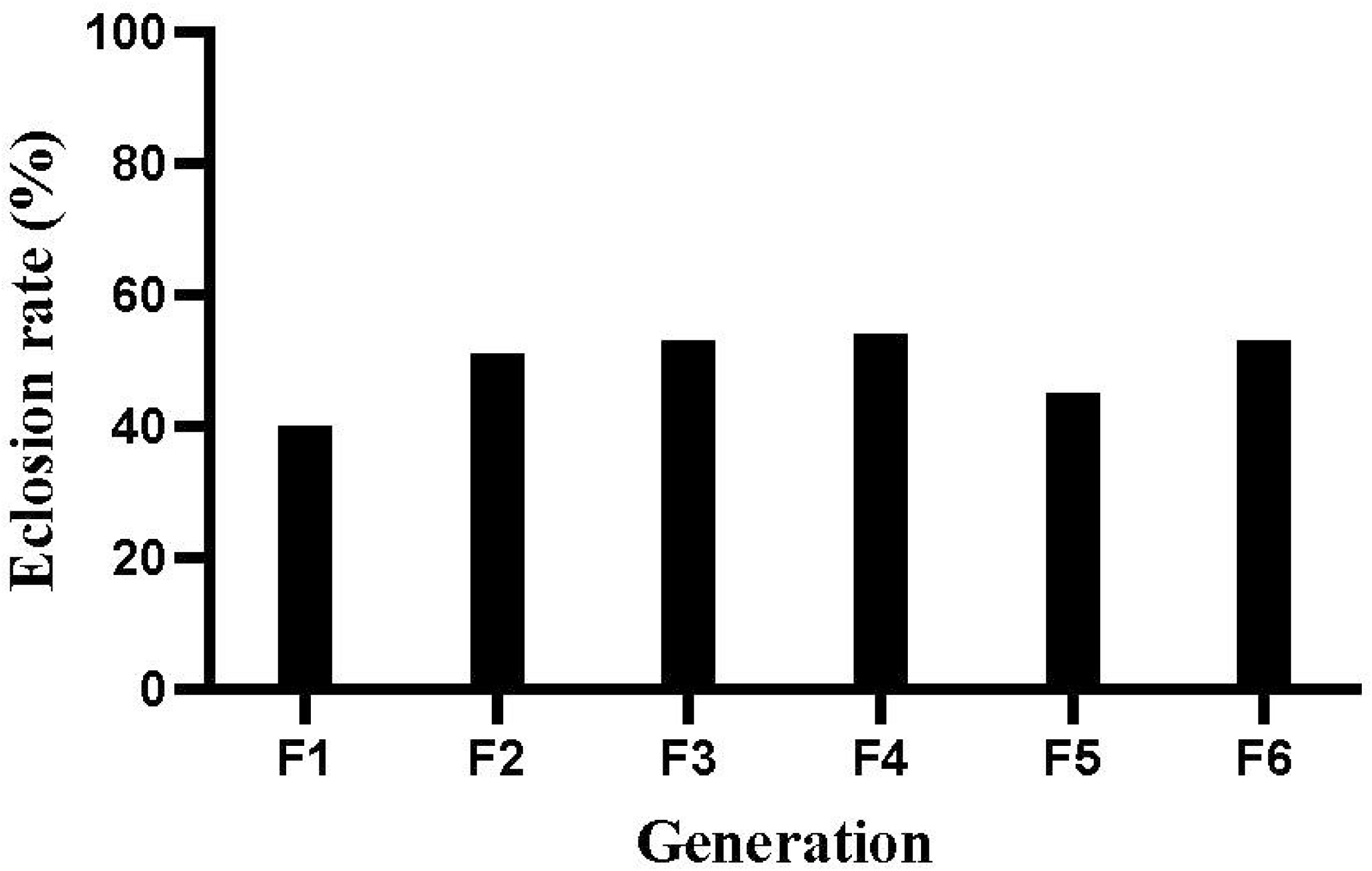
Egg eclosion rate at each generation. The number of L1 larvae hatching from each egg batch was determined by daily counting. L1 to L2 larvae hatching over the weekend were recorded on Mondays.

#### Larval and pupal development

The time between egg laying and first adult emergence was between 14 to 16 days under our insectary settings. The mortality rate of aquatic stages of *An. darlingi* was estimated as the ratio between the total number of adults and the total number of young larvae (L1 to L2) at each generation. As shown in Fig. 5, the aquatic stage mortality rate was generally lower than 25%, except in F4 where it reached 42%. The proportion of females stayed stable between 48 and 56% from F1 to F4, indicating that the sex ratio was not distorted from the expected 1:1 ratio (See Table 1).

**Figure 5:**
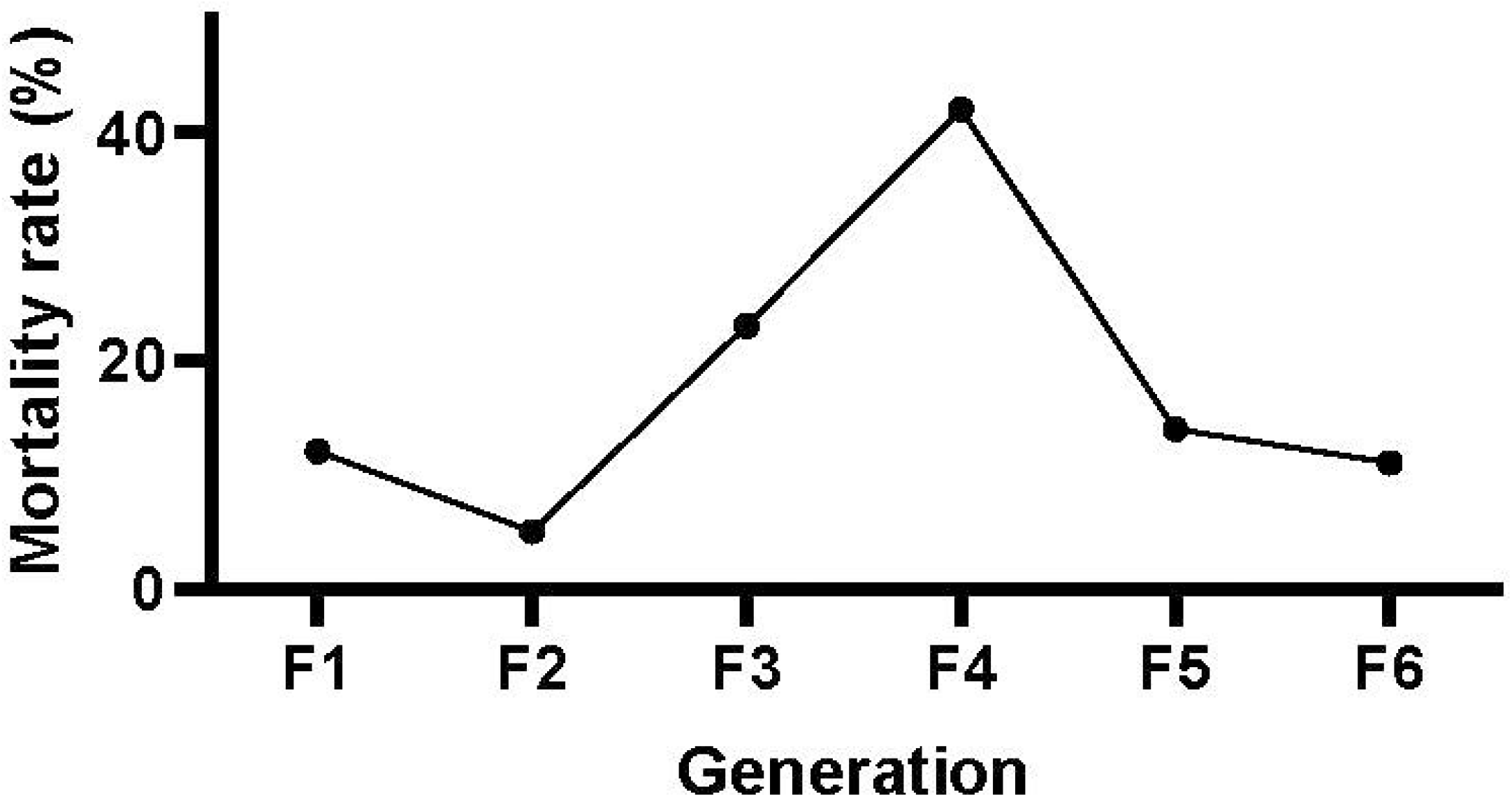
Larval survival success to adulthood across the first 6 generations of *An. darlingi*. The survival success is visualized from the mortality rate of larvae that did not reach adulthood, estimated from F1 to F6.

#### Monitoring the effect of the blue light flashes and shift in night temperature on free mating

To test whether the blue light flashes and the shift in night temperature were mandatory to maintain the colony as free mating after the first five generations, part of F5 adults, thereafter called F5’, were moved to a similar insectary without any light flashes and with constant temperature (26°C) producing the F6’, while the remaining F5 adults were maintained with blue light flashes and low night temperature producing the F6 generation. F7 and F7’ generations were obtained from F6 and F6’ respectively. Conversely, the next 2 generations from F7’ parents were submitted to light flashes while maintained at constant temperature (26°C). As presented in Table 2, we did not detect any significant differences in the egg production between F and F’ at each generation, whether or not normalizing data to the number of blood feedings each generation received (Sup data S2; Wilcoxon test on eggs/female – p=1; on egg/female/blood feed – p=1). This suggests that passed the fifth generation, neither light flashes nor a shift in night temperature is required for maintaining this free-mating *An. darlingi* colony.

**Table 2:**
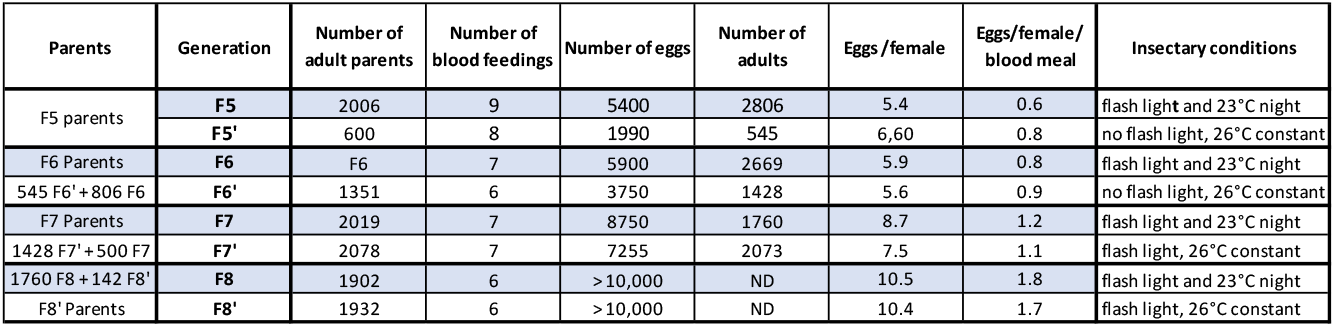
Effect of the blue light flashes and shift in night temperature on free mating of *An. darlingi*. At generations 5-8, the number of eggs/female is not affected by light flashes and temperature shifts (Wilcoxon test F vs F’–p=1).To keep similar numbers of adults between the F and F’ generations we mixed F and F’ parents in three instances (F6’, F7’ and F8).

| Parents | Generation | Number of adult parents | Number of blood feedings | Number of eggs | Number of adults | Eggs /female | Eggs/female/ blood meal | Insectary conditions |
| --- | --- | --- | --- | --- | --- | --- | --- | --- |
| F5 parents | F5 | 2006 | 9 | 5400 | 2806 | 5.4 | 0.6 | flash light and 23°C night |
|  | F5' | 600 | 8 | 1990 | 545 | 6,60 | 0.8 | no flash light, 26°C constant |
| F6 Parents | F6 | F6 | 7 | 5900 | 2669 | 5.9 | 0.8 | flash light and 23°C night |
| 545 F6' + 806 F6 | F6' | 1351 | 6 | 3750 | 1428 | 5.6 | 0.9 | no flash light, 26°C constant |
| F7 Parents | F7 | 2019 | 7 | 8750 | 1760 | 8.7 | 1.2 | flash light and 23°C night |
| 1428 F7' + 500 F7 | F7' | 2078 | 7 | 7255 | 2073 | 7.5 | 1.1 | flash light, 26°C constant |
| 1760 F8 + 142 F8' | F8 | 1902 | 6 | >10,000 | ND | 10.5 | 1.8 | flash light and 23°C night |
| F8' Parents | F8' | 1932 | 6 | >10,000 | ND | 10.4 | 1.7 | flash light, 26°C constant |

Some F13 eggs were sent to Institut Pasteur de la Guyane (Cayenne, French Guiana) to rear a duplicate colony. In both laboratories, generation F20 is currently ongoing.

### An. darlingi permissiveness to P. berghei and P. falciparum infection

We then tested whether this *An. darlingi* colony could be a valuable tool for vector competence studies. To this aim, *An. darlingi* and *An. stephensi* females were first infected in parallel with *P. berghei*. None of the 22 blood-fed *An. darlingi* females developed any oocyst whereas 75% of the blood-fed *An. stephensi* females (n=22) were infected with a mean oocyst load of 85.5 oocysts per infected female (Range 27-100; Fig. 6A). This is the first report revealing a possible refractoriness of *An. darlingi* to a rodent malaria parasite.

**Figure 6:**
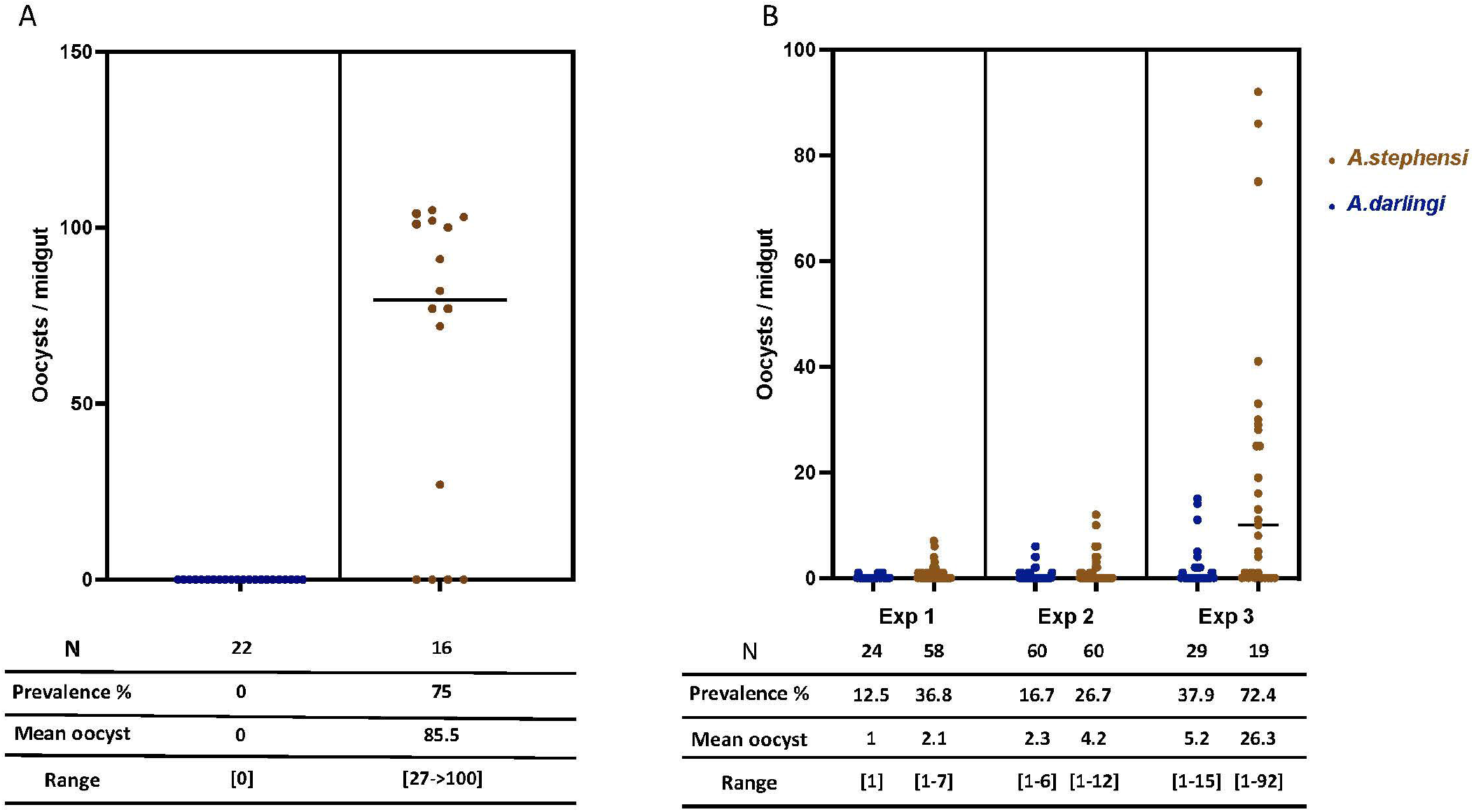
Infection of *An. darlingi* with *P. falciparum*. Five-day old *An. darlingi* females and *An. stephensi* (Sda 500) were starved overnight from sugar and fed on a *P. falciparum* NF54 gametocytes. Three experimental infections were performed using F9, F10 and F13 *An. darlingi* females, corresponding to Exp 1, 2 and 3 respectively. The horizontal bar on the graph represents the median of developing oocysts per mosquito midgut. Detailed data on prevalence and infection intensity are indicated below. Mean oocyst counts, and data range derive from infected midguts only.

Next, *An. darlingi* and *An. stephensi* females were infected with the same batch of *in vitro* produced *P. falciparum* gametocytes. Across 3 independent infection experiments, *An. darlingi* was found susceptible to *P. falciparum* NF54 (Fig. 6B). However, both infection prevalence and oocyst load were lower than the ones *o*f *An. stephensi*.

## Discussion

Here we report on the successful establishment of a free-mating *An. darlingi* colony starting from wild caught females in French Guiana. *An. darlingi* has a wide geographic distribution from central America to many countries in South America including the diverse regions of the amazon basin where this species was identified as a dominant malaria vector (in [1]). Based on various genetic analyses *An. darlingi* is composed of clearly isolated populations or clades [8, 24]. The present colony is likely representing a novel genetic resource of *An. darlingi.that* will offer additional values for a better characterization of the genetic diversity and vectorial competence of this major South America malaria vector.

The colony was established from the progeny of 34 wild caught females. This limited number of founders provided a starting pool of roughly 2,000 eggs. This number of founders is quite low compared to the numbers used for the most recently established *An. darlingi* colonies which were initiated from 135 up to 878 wild caught females [11, 17, 18].

To establish this colony, we developed a pragmatic approach by pooling information from our preliminary experiments and from published records on *An. darlingi* colonization since 1947. These data indicated that female insemination was a key parameter, and that forced mating did not allow fertilization, even though this technique was easy to perform with *An. darlingi* ([11, 17, 19], data not shown). We combined two main strategies. The first one capitalized on recent data obtained with *An. pseudopunctipennis* [19, 20] and *An. darlingi* [11] that suggest that mosquito copulation can be induced by providing light flashes just after dusk and decreasing night temperature as happening in the real mosquito life after dusk. The second strategy was to provide repeated blood meals to female mosquitoes up to 14 per generation over the first 6 generations. Providing light flashes after dusk as a copulation inducer was used for the recently established *An. darlingi* colonies from Peru and Brazil [11, 17, 18] while decreased night temperature was mentioned in only two reports [11, 18]. Our data reported in Table 2 show that neither light flashes nor night temperature shift are important past the 6^th^ generation. While we did not assess their role in the first six generations, we observed sexual activity as soon as light flashes started for the F1. Back to 1947, Bates observed “sexually excited males and mating” after exposure to lights of different colors [13]. He reported that mating occurred in the presence of a dim blue light when a limited number of mosquitoes (328 adults) were kept in a small cage, while mosquitoes did not mate without artificial light. In our insectary conditions, even if copulation was induced after dusk using light flashes, copulation was also observed in the daytime under dim light past the 10^th^ generation. Similarly, Villarreal-Treviño et al reported a very strong impact of the light-temperature combination to favor insemination at the 2^nd^ generation while light stimulation was not required anymore after the 10^th^ generation [11].

Because of limited space we chose middle size adult cages (47.5×47.5×47.5 cm) despite data suggesting that larger cages (60×60×60 cm) or even bigger (1.2m^2^x3m high) do better [11–14]. As recommended for *Anopheles funestus*, another difficult mosquito species to rear, and tested for *An. darlingi* [11], we used dark containers as egg laying device.

Following published data, mosquitoes were given their first blood meal on day 6 or day 7 post emergence to ensure that the copulation stimuli were efficient enough to produce a reasonable number of fertilized females at each generation. Mosquitoes were then fed almost every day on anesthetized rabbits or mice. Feeding mosquitoes on more than one type of animal blood was reported to favor female fecundity and we previously used this strategy to establish an *An. arabiensis* colony from Madagascar (Puchot & Bourgouin, unpublished).

As presented in Tables 1 & 2 and Sup Fig. S2 & S3, female fecundity (eggs/female) seems rather low at each generation. These numbers are in the same range as the ones obtained by Villarreal-Trevino *et al*. [11], using the same calculation method taking into account the entire female population rather than individual counts per fertilized female [17]. It is noticeable that the egg to female ratio increased at the 8^th^ generation while decreasing the number of blood meal provided. The number of blood meal has been reduced to 6-7 from the 9^th^ generation and to 3 since the 13^th^ generation. Meanwhile, we stopped counting the eggs as we had enough to proceed to the next generation.

Regarding the development time from egg to adult we obtained very similar results to published data in the range of 14 to 16 days [11, 12]. Noticeable was the asynchronous larval development from a same egg batch. Whether this was due to genetically different embryonic or larval development time or to the quantity and the type of the food provided to the larvae will need further investigation.

Importantly, we demonstrated that our free-mating *An.darlingi* colony is susceptible to *in vitro* produced gametocytes from *P. falciparum* NF54 originating from Africa. By comparison with concomitant *An. stephensi* control infections, the prevalence of infection was however slightly lower than the one obtained with F1 *An.darlingi* from Belize infected with the same *P. falciparum* strain [25].This difference may reflect an overall different genetic background between Belize and French Guiana *An.darlingi* populations or the selection of genotypes along the establishment of the colony from a limited number of founders. Interestingly, among the other species of th*e Nyssorhynchus subgenus, An. albimanus* is moderately refractory to *P. falciparum* NF54, whereas it is more permissive to *P. falciparum* 7G8, originating from Brazil [26]. By contrast *An. aquasalis* is refractory to both *P. falciparum* strains unless its immune system was affected by silencing the mosquito complement factor LRIM1 (Leucin-Rich-repeat-containing IMmune gene 1) or antibiotic treatment to remove their microbiota [27]. When challenged with *P. berghei*, our colony showed full refractoriness, as observed for *An. aquasalis* [27], suggesting that this parasite species is not a suitable model to study *An. darlingi*-*Plasmodium* interactions. Further investigation would be necessary to determine whether *P. berghei* is developing more slowly in *An. darlingi*, as previously shown in *An. albimanus* [28], and whether *Plasmodium yoelii* is a suitable rodent malaria model in *An. darlingi*, as demonstrated in *An. albimanus* [27].

In conclusion, this novel *An. darlingi* colony established from French Guiana mosquitoes offers an invaluable tool for a better understanding of *An. darlingi* genetic diversity, physiology and vector competence towards South America malaria parasites including *P. falciparum* and *P. vivax*.

## Author Contributions

Conceived the project and funding: CB, JBD, MG. Designed the experiments: NP, CB. Performed the experiments: NP, MTL, RC. Analyzed the data: NP, CB. Drafted the paper: CB, NP. Reviewed and edited the manuscript: NP, JBD, MG, CB.

## Acknowledgment

We thank members of the CEPIA platform for infecting our mosquitoes (*An. darlingi* and *An. stephensi*) with *P. berghei* and *P. falciparum*. Thanks to current and previous members of the Vectopole, especially Pascal Gaborit for technical help and to Anavaj Sakuntabhai for his constant support.

## References

1. Sinka, M., et al., A global map of dominant malaria vectors. Parasites & Vectors, 2012. 5(1): p. 69.

2. Baker, R.H., W.L. Drench, and J.B. Kitzmiller, Induced copulation in Anopheles mosquitoes. Mosquito News 1962. 22(1): p. 16–17.

3. Ow yang, C.K., F.L. St Maria, and R.H. Wharton, Maintenance of a laboratory colony of Anopheles maclalatus (Theobald) by artificial mating. Mosquito News, 1963. 23: p. 34–35.

4. Baker, R.H., Mating problems as related to the establishment and maintenance of laboratory colonies of mosquitos. Bull World Health Organ, 1964. 31: p. 467–8.

5. Amir, A., et al., Colonization of Anopheles cracens: a malaria vector of emerging importance. Parasites & Vectors, 2013. 6(1): p. 81.

6. Horosko, S., J.B.P. Lima, and M.B. Brandolini, Establishment of a free-mating colony of Anopheles albitarsis from Brazil. Journal of the American Mosquito Control Association, 1997. 13(1): p. 95–96.

7. Marinotti, O., What is in a name? Anopheles darlingi versus Nyssorhynchus darlingi. Trends in Parasitology, 2021.

8. Conn, J.E. and P.E. Ribolla, Chapter 5 - Ecology of Anopheles darlingi, the Primary Malaria Vector in the Americas and Current Nongenetic Methods of Vector Control, in Genetic Control of Malaria and Dengue, Z.N. Adelman, Editor. 2016, Academic Press: Boston. p. 81–102.

9. Yuval, B., Mating Systems Of Blood-Feeding Flies. Annual Review of Entomology, 2006. 51(1): p. 413–440.

10. Lounibos, L.P., D.C. Lima, and R. LourenÁo-de-Oliveira, Prompt mating of released Anopheles darlingi in western Amazonian Brazil. Journal of the American Mosquito Control Association, 1998. 14 2: p. 210–3.

11. Villarreal-Trevino, C., et al., Establishment of a free-mating, long-standing and highly productive laboratory colony of Anopheles darlingi from the Peruvian Amazon. Malar J, 2015. 14: p. 227.

12. Giglioli, G., Laboratory colony of Anopheles darlingi. J Natl Malar Soc, 1947. 6(3): p. 159–64.

13. Bates, M., The laboratory colonization of Anopheles darlingi. J Natl Malar Soc, 1947. 6(3): p. 155–8.

14. Americano Freire, S. and G. Faria, Rearing and some Data on the Biology of A. darlingi. Revista Brasileira de Biologia, 1947. 7(1): p. 57–66.

15. Corrêa, R.R., et al. Informe sobre uma colônia de Anopheles darlingi. in XVIII Congresso Brasileiro de Higiene. 1970. São Paulo, Brasil: Sociedade Brasileira de Higiene, Rio de Janeiro.

16. Buralli, G.M. and E.S. Bergo, [Maintenance of Anopheles darlingi Root, 1926 colony, in the laboratory]. Rev Inst Med Trop Sao Paulo, 1988. 30(3): p. 157–64.

17. Moreno, M., et al., Infection of laboratory-colonized Anopheles darlingi mosquitoes by Plasmodium vivax. Am J Trop Med Hyg, 2014. 90(4): p. 612–6.

18. Araujo, M.D.S., et al., Brazil’s first free-mating laboratory colony of Nyssorhynchus darlingi. Rev Soc Bras Med Trop, 2019. 52: p. e20190159.

19. Lardeux, F., et al., Laboratory colonization of Anopheles pseudopunctipennis (Diptera: Culicidae) without forced mating. C R Biol, 2007. 330(8): p. 571–5.

20. Villarreal, C., et al., Colonization of Anopheles pseudopunctipennis from Mexico. J Am Mosq Control Assoc, 1998. 14(4): p. 369–72.

21. Vezenegho, S.B., et al., Anopheles darlingi (Diptera: Culicidae) Dynamics in Relation to Meteorological Data in a Cattle Farm Located in the Coastal Region of French Guiana: Advantage of Mosquito Magnet Trap. Environ Entomol, 2015. 44(3): p. 454–62.

22. Nepomichene, T.N., et al., Efficient method for establishing F1 progeny from wild populations of Anopheles mosquitoes. Malaria Journal, 2017. 16(1): p. 21.

23. Mitri, C., et al., Density-dependent impact of the human malaria parasite Plasmodium falciparum gametocyte sex ratio on mosquito infection rates. Proc Biol Sci, 2009. 276(1673): p. 3721–6.

24. Prado, C.C., et al., Behavior and abundance of Anopheles darlingi in communities living in the Colombian Amazon riverside. PLOS ONE, 2019. 14(3): p. e0213335.

25. Grieco, J.P., et al., Comparative susceptibility of three species of Anopheles from Belize, Central America, to Plasmodium falciparum (NF-54). J Am Mosq Control Assoc, 2005. 21(3): p. 279–90.

26. Molina-Cruz, A., et al., Plasmodium evasion of mosquito immunity and global malaria transmission: The lock-and-key theory. Proc Natl Acad Sci U S A, 2015. 112(49): p. 15178–83.

27. Orfano, A.S., et al., Plasmodium yoelii nigeriensis (N67) Is a Robust Animal Model to Study Malaria Transmission by South American Anopheline Mosquitoes. PLOS ONE, 2016. 11(12): p. e0167178.

28. Frischknecht, F., et al., Using green fluorescent malaria parasites to screen for permissive vector mosquitoes. Malar J, 2006. 5: p. 23.

