## Supplemental Figure 1 for "Establishment of a colony of *Anopheles darlingi* from French Guiana for vector competence studies on malaria transmission"

**Supplementary figures**

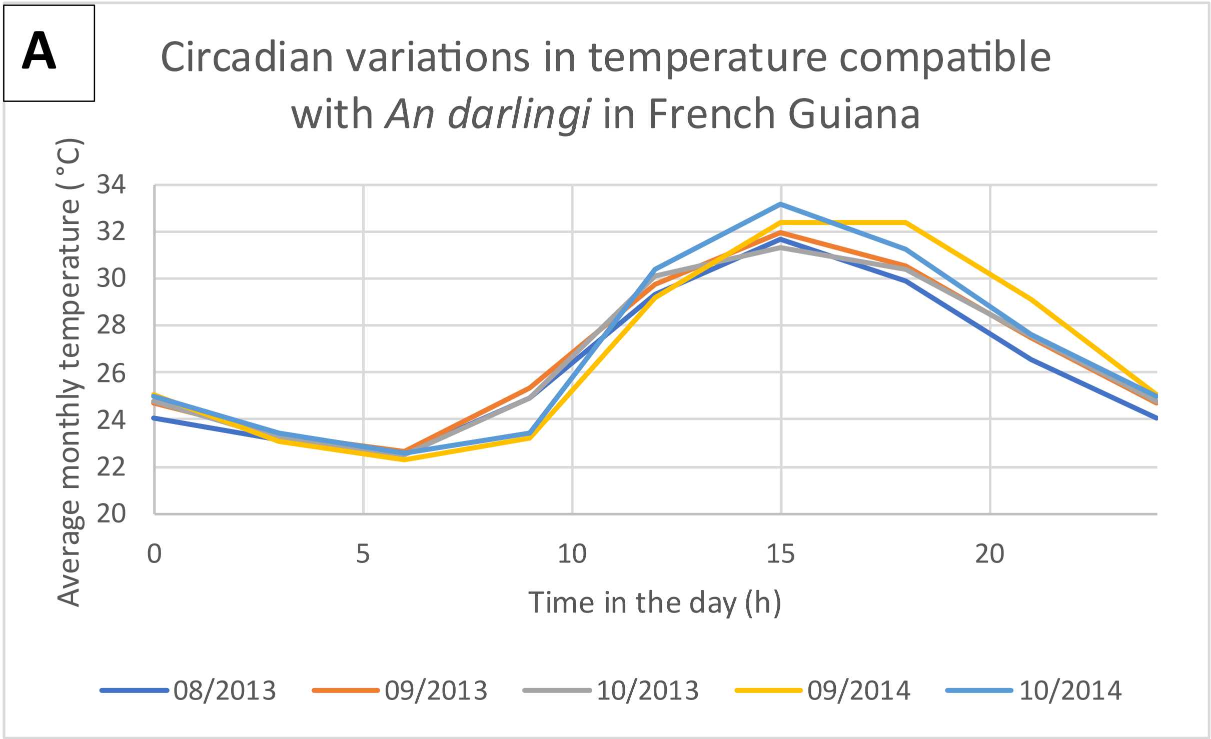


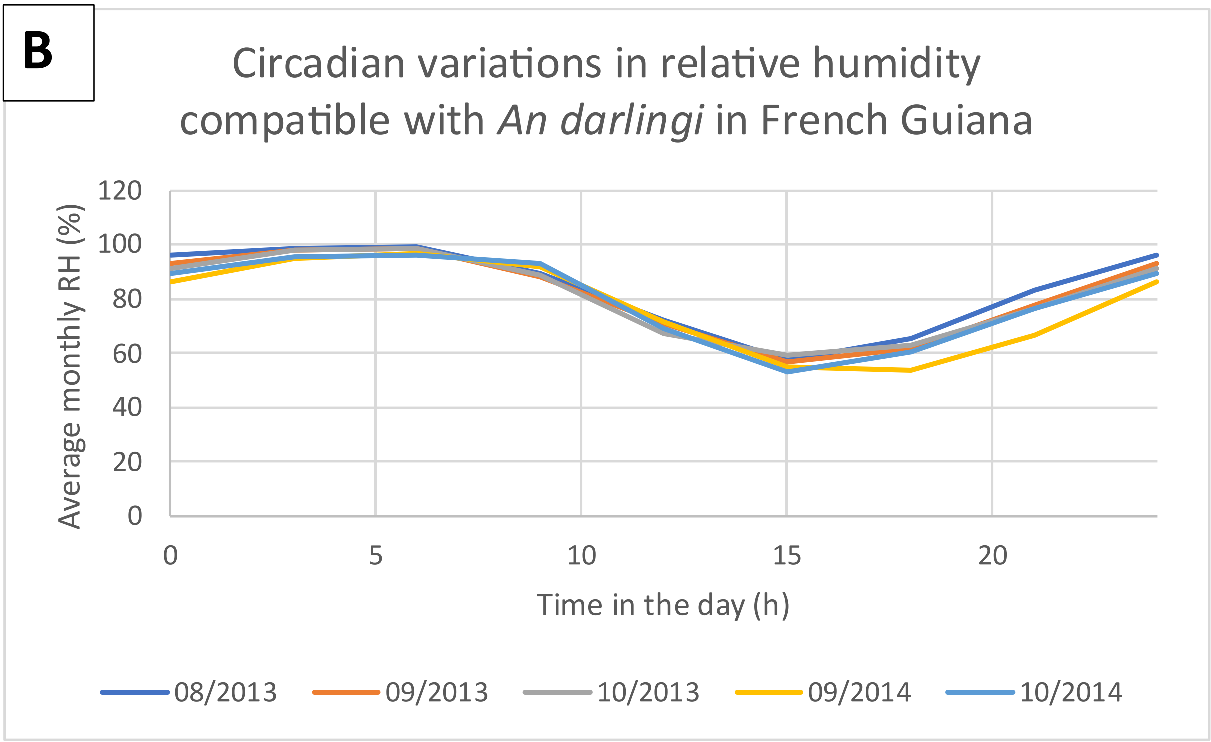


**Sup Figure S1** :Circadian variations in temperature (panel A) and relative humidity (panel B) in French Guiana. These circadian variations were observed in Saint Georges at specific months when *Anopheles darlingi* have been found in high quantity, as reported by Adde et al 2017 [1]. Data collected by Meteo France <2km apart from the mosquito collection site.

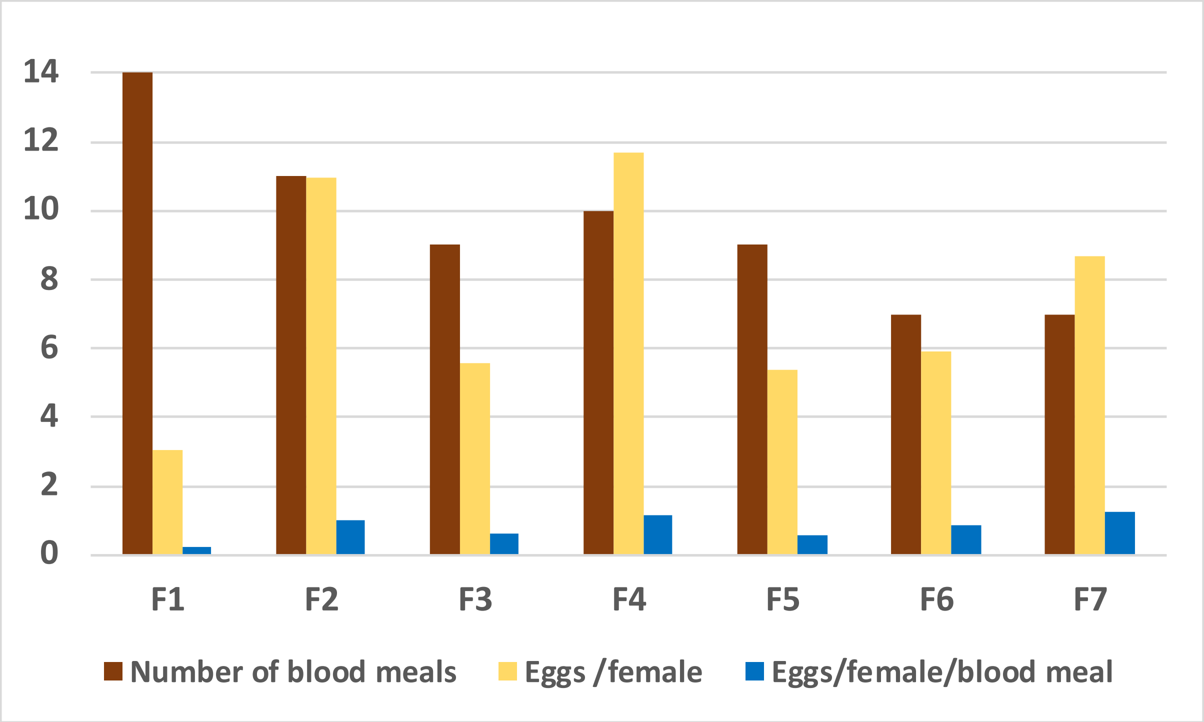


**Sup Figure S2** : Graphic view of the number of blood meals provided at each generation and the number of eggs per female and eggs per female and per blood meal. Data extracted from Table 1. See main text for estimating the egg to female ratio.

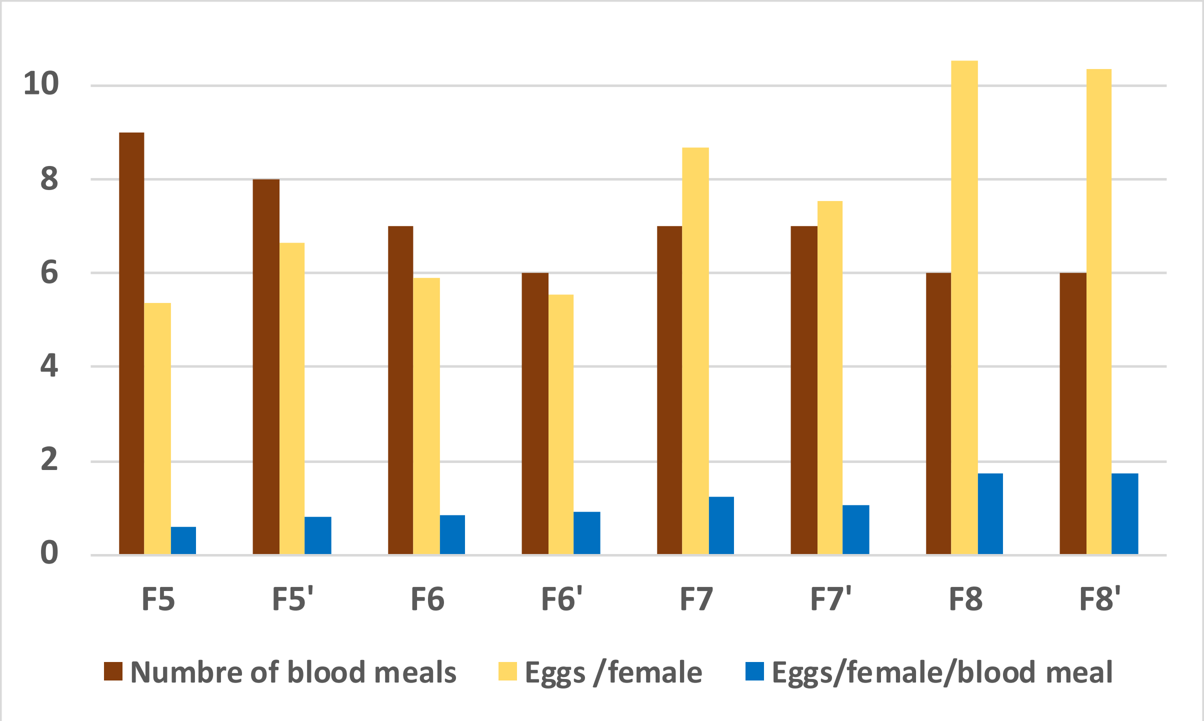


**Sup Figure S3**: Graphic view of the number of blood meals and the number of eggs per female exposed or not to the blue flashlight and to might temperature shift. Data extracted from Table 2. See main text for the description of the F & F’ generations.

**Reference:**


1. Adde, A., et al., *Spatial and Seasonal Dynamics of Anopheles Mosquitoes in Saint-Georges de l'Oyapock, French Guiana: Influence of Environmental Factors.* J Med Entomol, 2017. **54**(3): p. 597-605.
